# Vegetation and fires under climate change: a Mediterranean modelled case study from central Italy

**DOI:** 10.64898/2026.05.29.728691

**Authors:** Nicolò Perello, Gabriele Vissio, Pegah Aflakian, Guido Biondi, Mirko D’Andrea, Andrea Trucchia, Mara Baudena, Paolo Fiorucci

## Abstract

Wildfire regimes in Mediterranean landscapes are undergoing significant changes due to the combined effects of land-use transitions and climate change. In particular, land abandonment increased fuel availability, the expansion of the wildland–urban interface increased ignition frequency, while climate change increases the chances of fire-weather conditions and reduces vegetation recovery capacity. This study presents a modelling framework to investigate the coupled dynamics of fire and vegetation under different fire regimes scenarios, using a case study in central Italy (Monte Pisano). The approach integrates two cellular automata models for vegetation dynamics (Batllori et al., 2017) and for fire-spread (PROPAGATOR; Trucchia et al., 2020). The vegetation model represents succession among six functional classes, including grasslands, shrubs, and trees with different fire-response strategies (seeders and resprouters), while explicitly accounting for post-fire recovery processes. The model was calibrated for the area using historical fire perimeters and vegetation maps over 40 years. Fire spread is simulated probabilistically using PROPAGATOR, driven by fuel types, topography, and weather conditions. A stochastic coupling was implemented by sampling fuel classes from vegetation composition, and by feeding simulated burned areas back into the vegetation model, thus enabling dynamic fire-vegetation feedback. Future wildfire scenarios are constructed by linking ignition probability to fire-weather conditions derived from historical reanalysis data (1981–2023). Extreme fire events are defined based on thresholds of wind speed and fuel moisture, and their probability of occurrence is varied across scenarios to represent increasing climate-driven risk. Simulations are performed over a 100-year horizon starting from current vegetation conditions. Results show that, in the absence of fire, vegetation dynamics lead to dominance of late-successional, fire-resilient species (resprouters). This is particularly evident for low probabilities of extreme fire events, with fire impacts diminishing over time as landscapes become less flammable. However, increasing the frequency of extreme fire conditions resulted in persistent disturbance, maintaining higher proportions of shrubs and early successional vegetation, and sustaining elevated burned areas over time. Overall, the study shows that coupling fire spread and vegetation dynamics provides a useful framework for exploring long-term ecosystem trajectories under climate change. The results highlight the critical role of extreme fire events in shaping landscape resilience and suggest that future management strategies should account for fire–vegetation feedbacks to support more stable and less fire-prone ecosystems.

## 1. Introduction

For centuries, Mediterranean landscapes have been shaped by extensive land use, primarily driven by agricultural and pastoral activities. In recent decades, however, rapid land abandonment has transformed these landscapes into mosaics of highly flammable areas, many of which are transitioning toward ecosystems that are less vulnerable to fire. At present, high population density and the expansion of the wildland–urban interface have led to fire ignition frequencies that far exceed the time required for ecological succession to progress toward mature forest systems. In addition, ongoing climate change increases the likelihood of fire occurrence while simultaneously reducing the capacity of vegetation to recover, despite the many fire-adapted traits of Mediterranean species. As a result, management strategies aimed at preserving biodiversity across different successional stages, as well as maintaining the potential agricultural and forest productivity of these landscapes, face significant challenges - particularly in the context of current and future climate change.

This study aims to predict the future dynamics of fire regimes and ecosystems in the Mediterranean region under different climate change scenarios. To achieve this, we coupled a vegetation dynamics model - incorporating diverse post-fire response strategies (Batllori et al., 2017) - with a fire propagation model (Trucchia et al., 2020). Both models were calibrated for a representative area in central Italy, Monte Pisano, near Pisa.

## 2. Methodology

### 2.1. The vegetation model

The Batllori model (Batllori et al., 2017) describes the successional dynamics of four major plant functional types: grasses, shrubs, tree seeders (typically *Pinus* spp) and resprouters (typically sclerophyllous oaks). The trees are also divided in two demographic stages (young and mature), for a total of six classes. The model represents the proportion of vegetation cover at each site of a spatial lattice (conceptually thought to be a few tens of meters in size) and one-year timesteps. The successional dynamics is due to competition among species, modelled through replacement terms that depend on the interacting species pair. It also depends on seed availability: seeds are sourced from mature trees both locally and at the landscape scale. The model also represents the effect of fire events on species abundance, which can be summarised as a reversal in the succession process. The effect of fire on each species is characterised by their typical fire response (resprouters or seeders) and vegetation mortality rates are determined by species type and life stage. The young tree classes, representing seedlings and saplings, are more sensitive to fire than their mature counterparts, as they are unable to produce seeds and exhibit lower resprouting capacity. Unlike the original model, the approach presented here imposes historical data for forest fires (during calibration) and is coupled with PROPAGATOR (see below), thus we did not include the original ignition dynamics. We wrote the model code in Python following the model description in Batllori et al. (2017).

### 2.2. Coupling fire occurrence and spread to vegetation dynamics

PROPAGATOR is a stochastic cellular automata model for wildfire spreading. Given landscape description (i.e., topography and fuel types), ignitions, wind and fuel moisture conditions, the model produces a probabilistic wildfire spreading scenario for the imposed simulation time. The model adopts a stochastic logic of fire spreading between cells, influenced by the input information described above - by running different realizations of the stochastic process, burnt probability maps are produced at each time step, together with rate of spread and fire-line intensity maps. The model adopts customized fuel categories, adapted to the Mediterranean landscape: grassland, croplands, shrubs, conifers (corresponding to seeders in the vegetation model), broadleaves and low fire-risk forests (corresponding to resprouters). Each fuel type is associated with different rates of spread and intensity-related parameters, while the propagation process is guided by nominal propagation probabilities among the different fuel types.

In the present work, the PROPAGATOR model was used to run stochastic realizations of wildfire events in the study area, adopting as input the fuel map from the vegetational model and applying then the simulated wildfire burned areas as feedback into it. To associate to each cell a single fuel class, as required by PROPAGATOR, we adopted a stochastic approach: we randomly sampled a single fuel class for each realization of the propagation process between those present in the vegetation model cells, with a probability that was dependent on their cover/biomass.

The coupling between wildfire propagation simulator and vegetation model allowed for exploring different scenarios of future vegetational evolution at landscape level. An essential aspect of the statistical characterization of wildfire events is the identification and analysis of extreme fire events, which have the greatest impact on the territory and account for most of the burned area when an ignition occurs. The hypothesis adopted is that extreme fire events occur under extreme fire weather events. This characterization, in terms of both weather conditions and the probability of occurrence of extreme events, can be informed by the analysis of past events and by climate change scenarios for an area of interest. In this study we focused on past weather conditions analysis.

### 2.3. Application to real case study: the Monte Pisano

#### 2.3.1 Description of the study area

The Monte Pisano is a relatively small mountain system, with elevations below 1000 m, located in north-central Tuscany (Italy), with mean annual rainfall around 1200 mm and mean annual temperature around 15 °C, and surrounded by predominantly flat terrain. This area is characterized by vegetation of pines (mostly Maritime pine, *Pinus pinaster*), broadleaved evergreens (Holm oak, *Quercus ilex*), with deciduous trees such as Chestnuts (*Castanea sativa*) at the highest elevations and north facing slopes. The landscape also supports scattered shrublands. The area is also characterized by a long history of human influence, such as presence of olive groves, partly maintained and partly abandoned, and by recurrent wildfire events, making it an interesting case study, quite representative of mesic mediterranean conditions (Fig. 1). The area is supported by a substantial historical dataset, including land cover maps and wildfire records. Land cover maps provided by the Tuscany Region have been collected from regional geoportal (https://www.regione.toscana.it/-/geoscopio) in different years (1978, 2007, 2019) and used for calibrating the vegetation model. The maps were transformed from their original land cover values to the four land cover classes of the model. Fire perimeters available from Tuscany Region *(settore forestazione-ufficio AIB*) have been collected from the year 1970 to 2019 (Fig. 1). The area faced four main wildfires during the last 40 years (Tab. 1, burned areas greater than 400 ha), the main one occurred in 2018 with more than 1000 ha burned.

**Figure 1.**
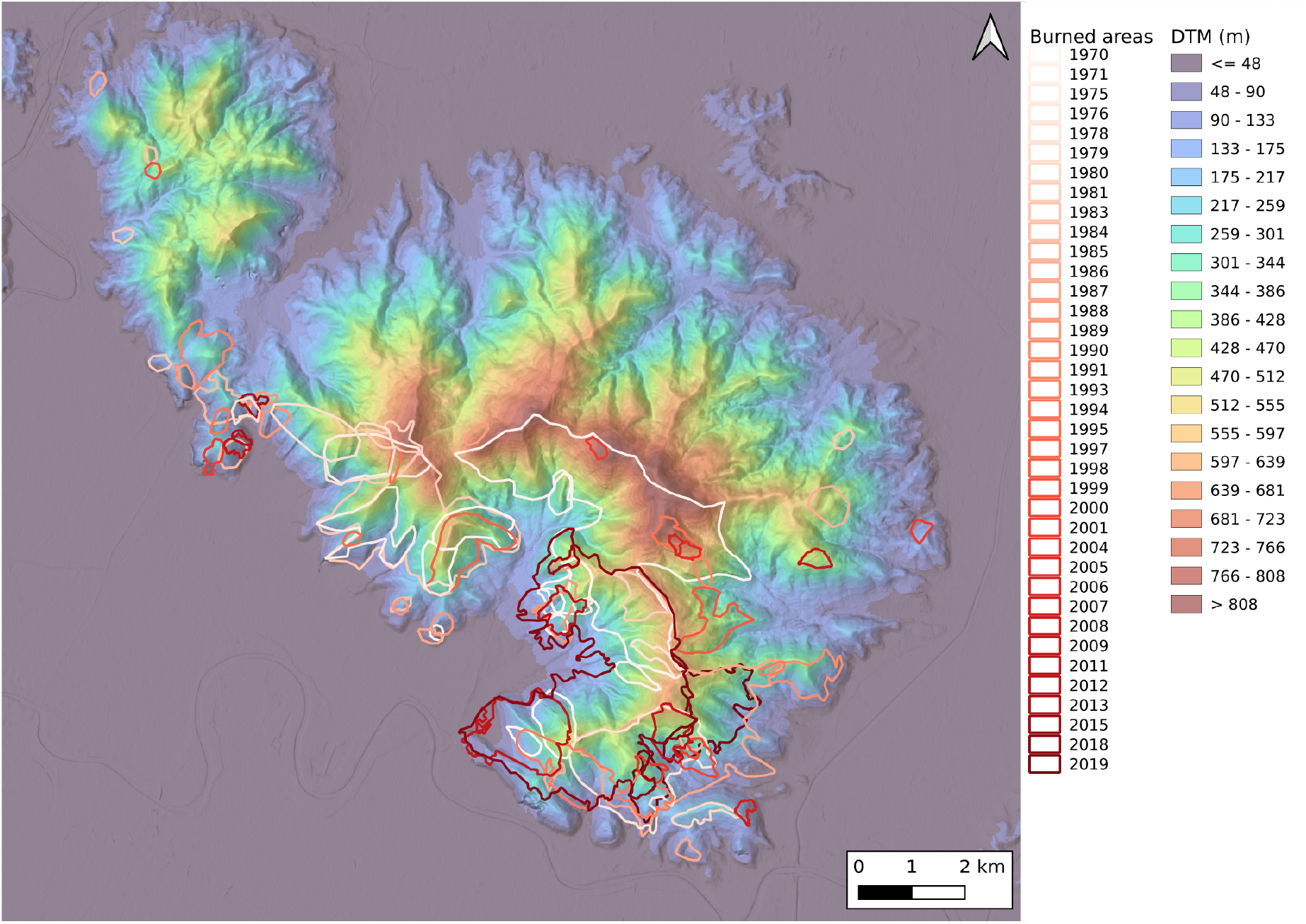
DEM map of the study area, including the perimeters of wildfires that occurred between 1970 and 2019.

**Table 1.** List of wildfires with burned area greater than 400 ha from 1970 to 2019.

| Year | Month | Burned area [ha] |
| --- | --- | --- |
| 1971 | 9 | 977 |
| 1983 | 7 | 495 |
| 1987 | 9 | 600 |
| 2018 | 9 | 1186 |

#### 2.3.2 Adapting the vegetation model to the Monte Pisano

In their original work, Batllori et al. (2017) assigned parameter values based on ecologically-informed assumptions rather than fitting them to observational data. We adopted a similar approach, applying ecological considerations to adapt some of their parameters to our study area. Regarding successional dynamics, the rate of change from grassland to shrubland was increased from 0.01 to 0.03, reflecting faster early-successional dynamics. Regarding post-fire dynamics, the fire-induced effective mortality of seeders was increased from 0.4 to 0.8 for young trees and from 0.25 to 0.7 for mature trees, in order to represent their weaker fire-response strategy relative to resprouters (Magnani et al., 2023).

We initiated a model run from the earliest forest map available (1978). We then run the model until the year 2019, including the information about when and whether cells burned. We performed a qualitative comparison between observed data and model output for 2007 and 2019 (Fig. 2), for the four forest classes. Analysing the differences between modeled and observed maps for the year 2019, we used confusion matrices (Fig. 3) to show the percentage of observed land cover assigned to each of the four classes by the model. We performed the analyses separately for the unburned cells, governed solely by successional dynamics, and the cells that burned at least once between 1978 and 2019, and therefore underwent also post-fire dynamics. Grasslands are systematically underrepresented across both dates, though it remains a minor class. The remaining three classes perform better overall, yet differently across the two conditions: broadleaf forest and conifers are well captured by the successional dynamics, whereas post-fire dynamics yield better results for shrubland, which is nonetheless overrepresented. Similar results were observed for the year 2007 (Fig. 2, confusion matrices are not shown).

**Figure 2.**
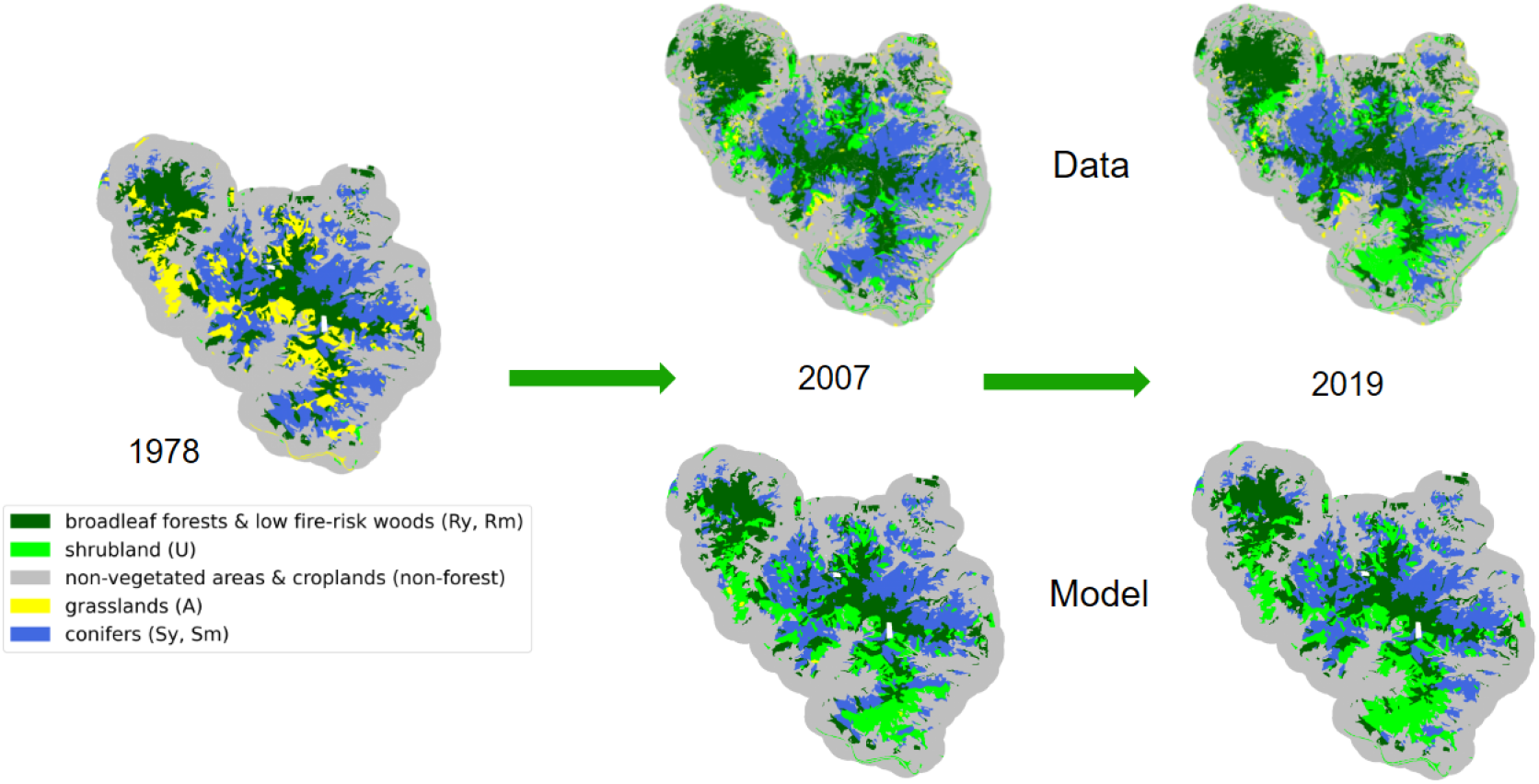
Comparison between modelled and observed land cover. Results for a run initialized in 1978. Top row: observed land cover; bottom row: simulated land cover.

**Figure 3.**
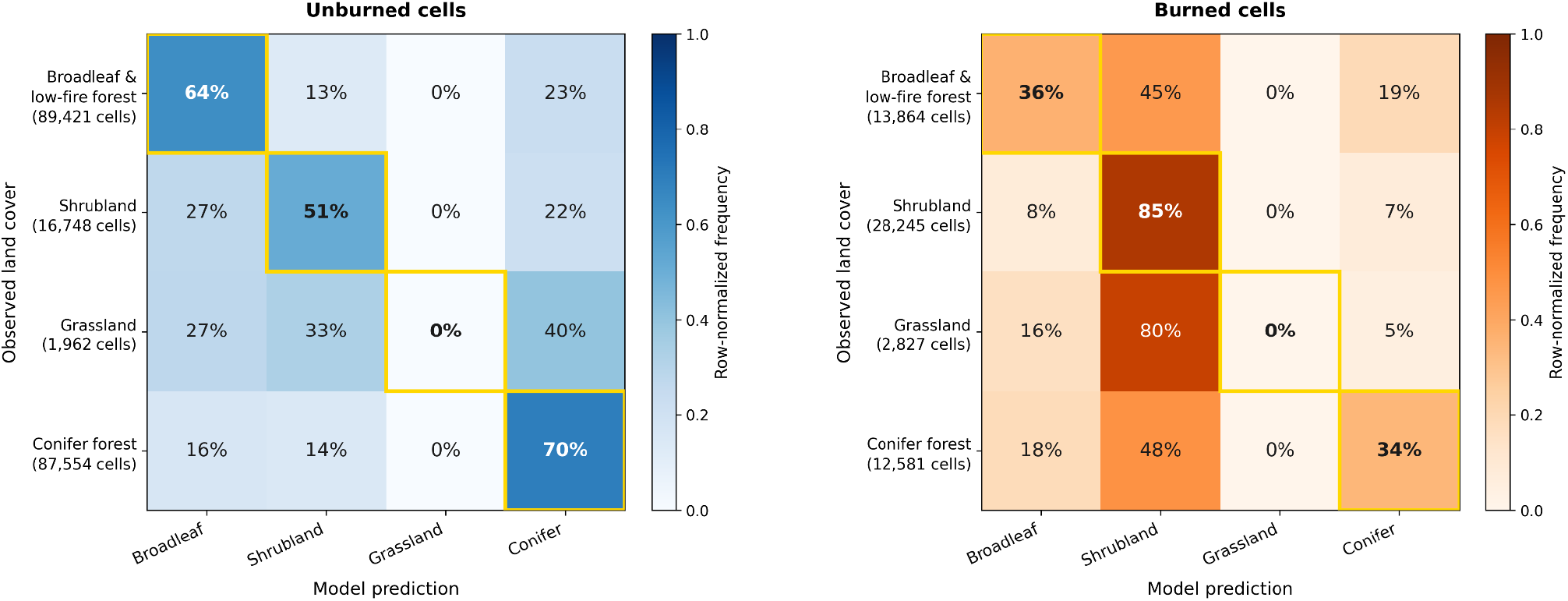
Confusion matrices for the year 2019 showing the percentage of observed land cover assigned to each vegetation class by the model (x-axis) in comparison to the observed forest map (y-axis). Left: unburned cells, governed by successional dynamics; right: cells burned at least once between 1978 and 2019, thus including post-fire dynamics.

#### 2.3.3 Definition of stochastic wildfire realizations and future scenarios

To generate future scenarios, we first needed to define a relationship between stochastic ignitions and fire-weather conditions. To do this, we analysed past wildfire events together with historical weather data derived from a high-resolution reanalysis dataset from 1981 to 2023 (Raffa et al., 2021). We defined extreme fire weather as conditions with wind speed greater than 5 m/s and relative humidity below 30% persisting for more than 6 hours. Based on these criteria, we counted the number of extreme weather events per year and found a slightly increasing trend in the study area (Fig. 4a). Monthly counts showed that March, April, and September were the most relevant months for extreme weather occurrence (Fig. 4b). However, when these results were compared with the seasonal distribution of wildfires in the area, particularly large fires exceeding 400 ha, March and April were not considered critical from a wildfire perspective. In those months, vegetation is generally not yet sufficiently dry to be highly flammable, whereas in September fuels are much more fire-prone after the summer period.

**Figure 4.**
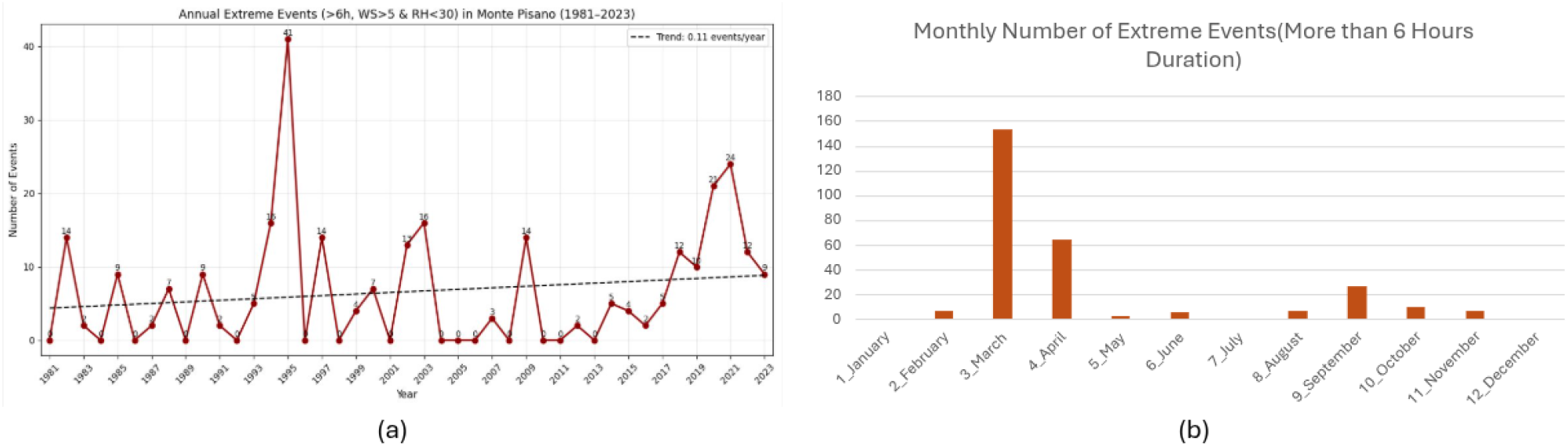
(a) Number of extreme fire weather conditions per year, from 1981 to 2023 (wind speed > 5m/s and relative humidity < 30% lasting for more than 6 hours); (b) number of extreme fire weather events per month, from 1981 to 2023.

Based on this analysis, we distinguished between normal fire-weather conditions, represented by wind speed of 5 km/h and fuel moisture of 15%, and extreme fire-weather conditions, defined as above and lasting 6 hours. The annual number of ignitions was then modelled using a normal distribution with a mean of 10 ignitions per year, while the probability that an ignition occurred under extreme fire-weather conditions varied depending on the scenario considered. As a baseline, we also tested a case with no extreme-fire ignitions. Given that 4 extreme fires were recorded in the last 40 years, the business-as-usual scenario was defined by assigning a 1% probability that an ignition would occur under extreme conditions. Two additional scenarios were then explored to represent increased extreme-fire likelihood: a medium-increase scenario with 5% probability, and a high-increase scenario with 10% probability.

## 3. Results and Discussion

The present manuscript reports some preliminary results.

With the coupled models, we simulated different scenarios of future trends for the next 100 years, starting from the last vegetation map available in the area (i.e. 2019). The scenarios differ by an increasing probability of extreme events (Fig. 5). In the Figure, the zero-condition (i.e. no fire events) is also shown as reference.

**Figure 5.**
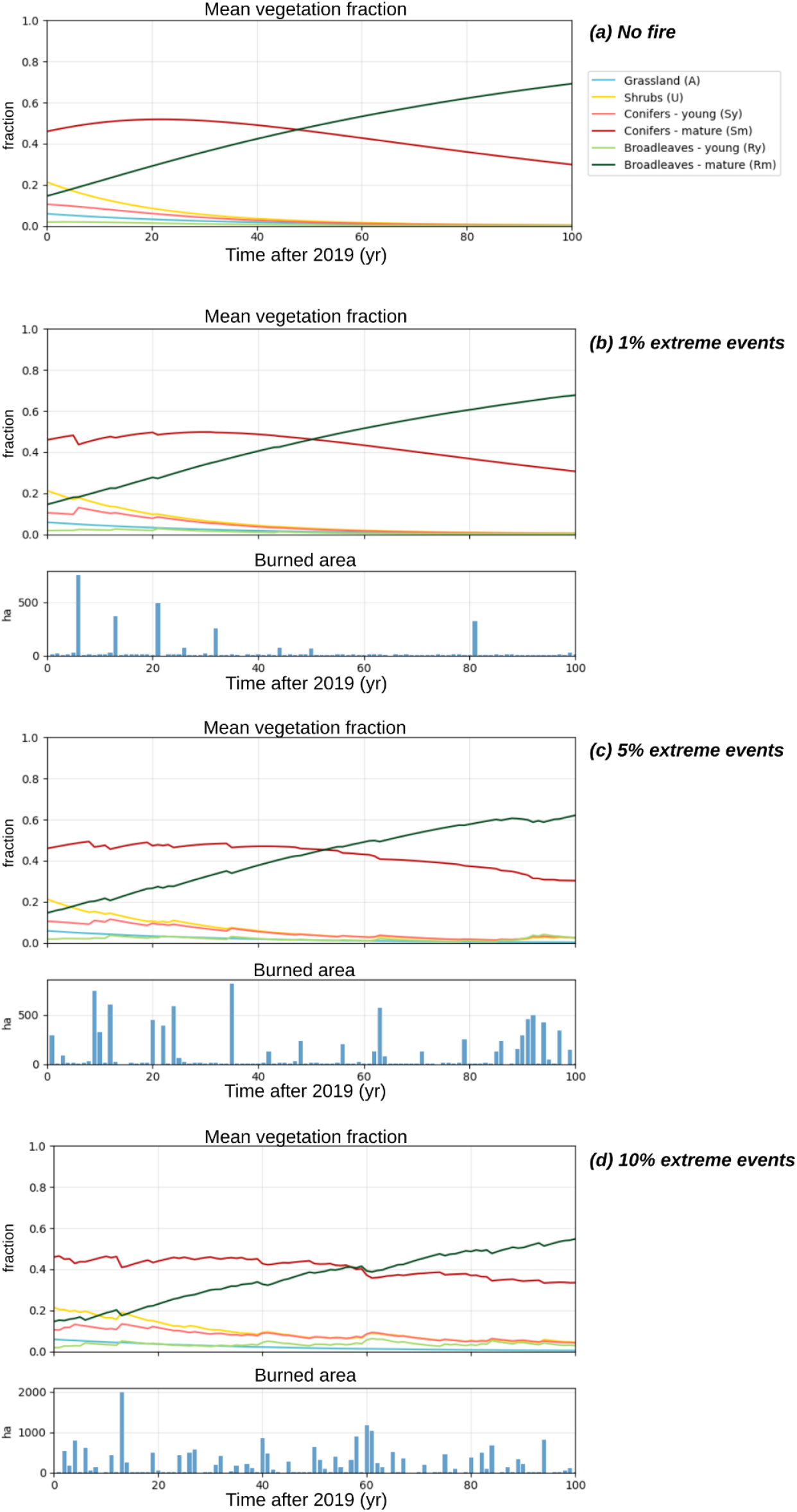
Mean fraction of each vegetation class as a function of time since 2019, averaged over the domain. Four different scenarios: (a) “no fire” scenario”; (b) 1% of extreme fire events scenario; (c) 5% of extreme fire events scenario; (d) 10% of extreme fire events scenario. In the bottom panels of (b-c-d), the burned area per year.

Vegetation dynamics as depicted by the Batllori model generally leads, over time, to the dominance of mature tree resprouters, i.e. the late successional species. This pattern is particularly evident, as expected, in the scenario without wildfires. Differences in the frequency of extreme events have a direct effect on the annual burned area and, consequently, on the progressively increasing influence of disturbance on successional dynamics, with fire representing the disturbance agent in this case.

As the frequency of extreme events increases, vegetation dynamics show progressively stronger disturbance-driven responses. In particular, the proportions of shrubs, and young seeders and resprouters classes tend to persist over time as a consequence of repeated fire occurrence (see for example *10% extreme events scenario*, Fig. 5d).

The probabilistic selection of the fuel map for the PROPAGATOR simulator, based on the proportions of vegetation classes, also affects the system dynamics. As the proportion of resprouters increases, because of both succession and resprouting abilities, the likelihood of assigning the PROPAGATOR broadleaved class also increases. This class is less conducive to fire spread than shrubs, grasslands, and conifers. Therefore, for the same probability of extreme events, the final burned area tends to decrease over time, along with the resulting impact of fire on vegetation - see for example *1% extreme events scenario*, Fig. 4, in which the effect of fire disturbances tends to be negligible after some years of simulation. In this sense, the system may reach a turning point at which landscape dynamics shift toward greater or lower resilience to extreme events. Conversely, when the frequency of extreme events increases, the burned area remains consistently high over time, owing to the persistence of a sufficient proportion of grassland, shrubland, and conifer-dominated areas, which are more conducive to fire spread.

## 4. Conclusions and Future Work

Coupling a vegetation model with a fire spread model, together with a probabilistic characterization of wildfire occurrence and fire weather conditions, can provide a useful framework for exploring the evolution of fire regimes and ecosystems under climate change. This approach could be extended to investigate management scenarios aimed at supporting the development of more resilient landscapes.

In this study, the Monte Pisano area was used as a case study, and past event analyses were employed to calibrate the model and develop future scenarios. To strengthen the robustness of the framework, future developments could incorporate scenarios with a gradual increase in the frequency of extreme events over time, as well as a more comprehensive probabilistic characterization of such events, in order to explore future scenarios more effectively. In addition, given the possibility of simulating fire behavior characteristics such as potential fire intensity and crown fire effects, stronger feedbacks between fire and vegetation dynamics could be included. Finally, the exploration of management scenarios represents a particularly relevant direction for future work, with the aim of enhancing the model as a practical tool for scenario analysis and decision support.

## 5. Acknowledgements

MB, PF, and GV were supported by the Italian National Biodiversity Future Center (NBFC), National Recovery and Resilience Plan (NRRP), Mission 4 Component 2 Investment 1.4 of the Italian Ministry of University and Research; funded by the European Union—NextGenerationEU (Project code CN00000033). The research activities described in this paper have been partially funded by MED-Star2 project - EU Program Interreg Italia-Francia Marittimo 2021-2027. This research was partially funded by the Italian Civil Protection Department Presidency of the Council of Ministers, through the convention with CIMA Research Foundation, for the development of knowledge, methodologies, technologies, and training, useful for the implementation of national systems of monitoring, prevention, and surveillance.

The authors acknowledge Antonello Provenzale, Fasma Diele and Carmela Marangi for discussions and Fasma Diele also for providing an earlier version of the vegetation model code.

## Notes

### Competing Interest Statement

The authors have declared no competing interest.

